# Bit-Reproducible Phylogenetic Tree Inference under Varying Core-Counts via Reproducible Parallel Reduction Operators

**DOI:** 10.1101/2025.06.02.656320

**Authors:** Christoph Stelz, Lukas Hübner, Alexandros Stamatakis

## Abstract

**Motivation:** Phylogenetic trees describe the evolutionary history among biological species based on their genomic data. Maximum Likelihood (ML) based phylogenetic inference tools search for the tree and evolutionary model that best explain the observed genomic data. Given the independence of likelihood score calculations between different genomic sites, parallel computation is commonly deployed. This is followed by a parallel summation over the per-site scores to obtain the overall likelihood score of the tree. However, basic arithmetic operations on IEEE 754 floating-point numbers, such as addition and multiplication, inherently introduce rounding errors. Consequently, the order by which floating-point operations are executed affects the exact resulting likelihood value since these operations are not associative. Moreover, parallel reduction algorithms in numerical codes re-associate operations as a function of the core count and cluster network topology, inducing different round-off errors. These low-level deviations *can* cause heuristic searches to diverge and induce high-level result discrepancies (e.g., yield topologically distinct phylogenies). This effect has also been observed in multiple scientific fields, beyond phylogenetics.

**Results:** We observe that varying the degree of parallelism results in diverging phylogenetic tree searches (high level results) for over 31% out of 10 130 empirical datasets. More importantly, 8% of these diverging datasets yield trees that are statistically significantly worse than the best known ML tree for the dataset (AU-test, *p <* 0.05). To alleviate this, we develop a variant of the widely used phylogenetic inference tool RAxML-NG, which does yield bit-reproducible results under varying core-counts, with a slowdown of only 0 to 12.7% (median 0.8%) on up to 768 cores. We further introduce the ReproRed reduction algorithm, which yields bit-identical results under varying core-counts, by maintaining a fixed operation order that is independent of the communication pattern. ReproRed is thus applicable to all associative reduction operations – in contrast to competitors, which are confined to summation. Our ReproRed reduction algorithm only exchanges the theoretical minimum number of messages, overlaps communication with computation, and utilizes fast base-cases for local reductions. ReproRed is able to all-reduce (via a subsequent broadcast) 4.1 *×* 10^6^ operands across 48 to 768 cores in 19.7 to 48.61 µs, thereby exhibiting a slowdown of 13 to 93% over a non-reproducible all-reduce algorithm. ReproRed outperforms the state-of-the-art reproducible all-reduction algorithm ReproBLAS (offers summation only) beyond 10 000 elements per core. In summary, we re-assess non-reproducibility in parallel phylogenetic inference, present the first bit-reproducible parallel phylogenetic inference tool, as well as introduce a general algorithm and open-source code for conducting reproducible associative parallel reduction operations.

**Availability and Implementation:** ReproRed: https://doi.org/10.5281/zenodo.15004918 (LGPL) — Reproducible RAxML-NG version https://doi.org/10.5281/zenodo.15017407 (GPL)

**Supplementary information:** https://doi.org/10.5281/zenodo.15524754

**Funding:** This project received funding from the European Union via European Research Council (ERC) Horizon 2020 research and innovation grant No. 882500 and via the EU ERA Chair (HORIZON-WIDERA-2022-TALENTS-01: 2023-2028) program grant No. 101087081 (Comp-Biodiv-GR).

## 1. Introduction

Next to disseminating results, scientific publications also aim to convince the reader of their validity [24]. While reproducibility is crucial for validating scientific claims [20], practical attempts to reproduce computational findings frequently fail, for example, in climate and weather modeling [10], power grid analysis [35], or phylogenetic tree inference [8, 30].

*Phylogenetic trees* describe the shared evolutionary history among related biological species based on their genomic data. Phylogenetic trees are commonly inferred via Maximum Likelihood (ML) based methods that search for the tree and evolutionary model that best explain the observed genomic data [11]. For this, the currently best-known tree is iteratively being improved by proposing topologically similar trees, and numerically optimizing the branch lengths and evolutionary model to assess if a proposed tree improves upon the score of the current best-known tree. To score a tree, statistical models of evolution are used to compute per-site *likelihood* scores. Given the independence of likelihood calculations between sites, phylogenetic inference tools are commonly parallelized over sites. Subsequently, summing over the per-site scores yields the overall tree score. If a proposed topology has a better likelihood score, we continue optimization from this new tree.

However, this iterative improvement of the best-known tree is known to be numerically unstable under varying CPU core counts [8, 29]. Even if an identical binary is re-executed on the same hardware but using a distinct number of threads/MPI processes, we can already observe diverging tree searches.

We define *bit-reproducibility* as a program producing bit-wise identical results when executed twice on the same input data and under the same settings. While tools such as containerization exist to archive the software environment and experimental settings [4], hardware is substantially more difficult to archive [20]. In order to reproduce results even without access to the hardware originally used, we thus require algorithms that yield bit-reproducible results on a plethora of hardware environments, including different CPU types and varying parallelization levels.

### 1.1 Contribution and Outline

We investigate the bit-reproducibility of phylogenetic tree inference across 10 130 empirical datasets under varying core counts and distinct SIMD parallelization kernels. Our findings qualitatively support those by Shen *et al*. [30]. We find that 31% of the tree searches diverge; with 8% (out of those 31%) yielding trees that are statistically significantly worse than the best known tree for the dataset (AU-test, *p <* 0.05). To address this issue, we introduce a bit-reproducible version of the phylogenetic tree inference tool RAxML-NG [21]. This bit-reproducible variant only exhibits a slowdown of 0 to 12.7% (median 0.8%) compared to the non-reproducible reference on 18 large empirical datasets when using up to 768 cores.

Further, we introduce the distributed memory parallel reduction algorithm ReproRed, which enables bit-reproducible reduction operations for arbitrary associative operators. This versatility sets ReproRed apart from competitors that are confined to summation (Section 3). It also allowed us to integrate ReproRed into the open-source MPI-wrapper library KaMPIng. ReproRed only exchanges the theoretical minimum number of messages, overlaps communication with computation, and utilizes a fast base-case for conducting local reductions. ReproRed performs an all-reduction (via a subsequent broadcast) on 4.1 *×* 10^6^ operands across 48 to 768 cores in 19.7 to 48.61 µs, thereby yielding in a slowdown of 13 to 93% over a non-reproducible all-reduce algorithm and 8 to 25% over a non-reproducible Reduce-Bcast algorithm. ReproRed outperforms the state-of-the-art reproducible all-reduction algorithm, ReproBLAS (summation only) for more than 10 000 elements per PE.

The remainder of our paper is organized as follows: In Section 2, we provide the necessary background and definitions for the remainder of our work. In Section 3, we discuss related work, outline the design of our operation-agnostic, bit-reproducible parallel reduction algorithm ReproRed in Section 4, and describe its efficient implementation Section 4.2. Next, in Section 5, we detail the bit-reproducible variant of the phylogenetic search tool RAxML-NG. In Section 6, we describe our experimental setup, and subsequently present our large-scale study on the reproducibility of phylogenetic tree searches in Section 6.1. Section 6.4 provides an empirical evaluation of ReproRed’s runtime compared to the state-of-the-art bit-reproducible (all-)reduction algorithm ReproBLAS. Following this, we assess the runtime penalty induced to RAxML-NG by using bit-reproducible reduction (Section 6.5), summarize our findings, and discuss directions of future research (Section 7).

## 2. Preliminaries

### 2.1 Parallel Reduction and All-Reduction

Given an input array of elements *E* = [*e*_0_, *e*_1_, …, *e*_*n−*1_], distributed across *p* Processing Elements (PEs; e.g., processes or threads), we desire to compute *r* = *e*_0_ ⊕*e*_1_ ⊕…⊕*e*_*n−*1_, where ⊕ denotes a binary, associative operation (e.g., summation or multiplication). In a distributed *reduction*, we return the result *r* to a single PE, while in a distributed *all-reduction*, we return *r* to all PEs. Conceptually, an all-reduction therefore comprises a reduction that is followed by a broadcast. However, this approach results in a latency of 2 *·* log_2_(*p*) messages: log_2_(*p*) messages for the reduction phase and log_2_(*p*) messages for the broadcast phase [28, ch. 13.2.1]. Dedicated all-reduction algorithms, such as recursive doubling which is implemented in MPICH [3], exhibit a latency of only log_2_(*p*) messages.

Next, we need to define a binomial tree: The binomial tree *B*_0_ of order 0 comprises a single node. A binomial tree *B*_*k*_ of order *k* entails a root node whose children are the roots of the binomial trees of order *k −* 1, *k −* 2, …, 2, 1, 0. Further, we define a generalized binomial tree *B*^*m*^, where the root of a binomial tree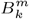has *m −* 1 copies of each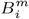 for *i* = *k −* 1, *k −* 2, …, 2, 1, 0 as children. Thus, in this notation, a canonical binomial tree is a *B*^2^. In a binomial tree reduction, PEs send intermediate results along the edges of a binomial tree towards the root and also reduce the values in each step. When we only have but a few elements per PE, a binomial tree reduction is optimal with respect to message count, size, and arithmetic operations – and thus, also regarding runtime [28, ch. 13.1.1].

KaMPIng [34] is a high-level C^**++**^ wrapper for MPI, which includes the reduction and all-reduction collective operations. The user can provide arbitrary associative reduction operations to KaMPIng’s `reduce` and `allreduce` functions. Note that associativity is a necessary requirement for a *parallel* reduction.

### 2.2 Floating-Point Math is Non-Associative

Fundamental arithmetic operations, such as additions or multiplications, on IEEE 754 [17] floating-point numbers induce rounding errors [13]. Thus, the compiler’s choice of CPU instructions (e.g.,, Fused-Multiply-and-Add [18, ch. 14.5.2]), which is often based on the available CPU type, influences the magnitude and propagation of these rounding errors. In addition, inconsistent floating-point related CPU settings can also cause different rounding errors. Examples include the IEEE 754 rounding mode [18, ch. 4.8.4], denormalized floating-point numbers [18, ch. 10.2.3.4], floating-point exceptions [7], and x87 register precision [18, ch. 8]. Henceforth, we therefore assume that the exact same instructions are used in each program execution. This can be achieved, for instance, by archiving the compiled binary or by passing the respective reproducibility flags to the compiler [7].

### 2.3 Non-Reproducibility of Distributed Reduction

In common distributed-memory parallel reduction algorithms, the PE-count influences the operation order and thus, the end-result [23, 32] (Figure 1). This is because the data are often distributed across all PEs in parallel algorithms, with each PE contributing a partial result to the reduction. Changing the parallelization level induces a different data distribution and communication pattern, which in turn alters the order of reduction operations. In addition, topology-aware reduction algorithms, such as in MPICH, adjust the reduction tree topology to accommodate for the heterogeneous connectivity among PEs [3], and thus yield non-reproducible results even under a constant PE-count.

## 3. Related Work

### 3.1 Non-Reproducible Phylogenetic Inferences

Darriba *et al*. [8] first reported that ML phylogenetic tree searches can diverge due to floating-point inaccuracies. Following this observation, Shen *et al*. [30] systematically analyzed 3515 single-gene datasets. They observed that variations in the CPU model, SIMD instruction set being used, and the number of threads, yielded different phylogenetic trees for 9 to 18% of empirical single-gene datasets. Furthermore, Shen *et al*. observed that 8.6% of the non-reproducible phylogenetic trees found by IQ-TREE and 25.21% of the non-reproducible phylogenetic trees found by RAxML-NG were statistically significantly worse than the respective best-known tree (*p <* 0.05; AU-test [31]).

### 3.2 Reproducible Reduction

(All-)reduction algorithms that yield bit-identical results under varying PE-counts have been studied for both, shared-memory [35] and distributed-memory [5, 16] systems.

Villa *et al*. [35] developed a bit-reproducible reduction algorithm on a shared memory Cray XMT machine and utilized it to sum over floating-point numbers on up to 16 PEs. Their algorithm ensures bit-reproducibility by fixing the reduction operation order independently of the PE-count, thereby supporting arbitrary associative reduction operations. Their *reduction* tree is based on a binomial tree, however, they accumulate *k >* 2 instead of 2 values sequentially at each inner node, which reduces the required tree height. Villa *et al*.’s algorithms is not open-source and is designed for a shared-memory Cray system (which we do not have access to), yielding a comparison to our distributed-memory algorithm infeasible.

For distributed-memory machines, one has to consider explicit communication between PEs as well as scheduling the reduction operations to PEs. Although the MPI standard encourages bit-reproducible reduction implementations [25, ch. 6.9], existing MPI implementations prioritize performance over reproducibility [3]. While IntelMPI and MPICH also offer reproducible reduction [19, 3], they both require an identical PE-count between runs. Further, Balaji and Kimpe [3] claim that reproducible reduction cannot utilize knowledge about the concrete HPC topology for improving communication efficiency. However, we separate the communication pattern from the computation of the reduction and thereby demonstrate that this is not necessarily the case.

Numerous bit-reproducible parallel *summation* algorithms exists [10, 5, 6, 2, 23, 1]. These algorithms utilize variants of Kahan’s compensated summation [32], which accumulates errors from each pair-wise summation in a separate variable. Thereby these algorithms effectively increase the number of significant bits. Demmel *et al*. [9] extend this concept to multiple variables, where each variable successively accumulates smaller parts of the error [32]. However, these approaches are summation-specific and do not generalize to other reduction operations. Further, most of these approaches require a specific data representation and have a single target hardware architecture, which complicates code maintenance [32].

The above algorithms have been implemented in three highly-optimized BLAS libraries: ReproBLAS [1], RareBLAS [5], and ExBLAS [16]. However, RareBLAS [5] targets parallel shared-memory machines, yielding it inapplicable to our distributed-memory context. Further, Lei *et al*. [22] show that ReproBLAS is consistently faster than ExBLAS. Consequently, we omit ExBLAS and only consider ReproBLAS as the state-of-the-art implementation for evaluating our ReproRed algorithm.

### 3.3 Further Baselines

The *Gather-Bcast* algorithm initially gathers all input data elements on a single root PE, which subsequently reduces the data elements in a fixed order, for instance, from left to right. The root PE then broadcasts the reduced result to all other PEs. The MPI standard recommends this algorithm for obtaining bit-reproducible results under varying PE-counts [25, ch 6.9]. However, this approach incurs a bottleneck communication volume of *p −* 1 messages and *n* received elements at the root PE, where *p* is the number of PEs, and *n* is the number of input data elements. Further, this algorithm lacks parallelism, resulting in a runtime of *O*(*n*). Moreover, the root PE requires sufficient memory to hold *all* input data elements.

We also compare ReproRed against two *non-reproducible* baselines: A PE-local reduction using C^**++**^’s `std::reduce` followed by either IntelMPI’s `MPI Allreduce`, or a `MPI Reduce` and a subsequent `MPI Bcast`. We denote these algorithms as *Allreduce* and *Reduce-Bcast*, respectively. In our experiments, IntelMPI selects the “best” (all-)reduction algorithm by using yet undisclosed heuristics. We attempted to re-configure IntelMPI to use a binomial tree broadcast, binomial tree reduction, and recursive doubling all-reduction by adjusting the respective `I MPI ADJUST` variables. However, this resulted in performance degradation, possibly because of the loss of topology-aware optimizations. We therefore perform all benchmarks under IntelMPI’s default settings.

## 4. Bit-Reproducible Parallel Reduction

The only approach for ensuring bit-reproducibility for arbitrary (associative) reduction operations, is to maintain a fixed reduction operation order [2]. To this end, we orchestrate the reduction operations via a binary tree with the input data elements located at its leaves. This *pair-wise reduction* also decreases the overall rounding errors [15]. Further, we orchestrate communication via a binomial tree, which is theoretically optimal for small message sizes [28, ch. 13.9], and used, for instance, by MPICH [3]. Note that this design implies that PEs can generally not reduce all their local elements into a single intermediate result, but are required to send the intermediate results of multiple subtrees to their parent in the communication tree instead. However, we expect the startup overhead for each message to exceed the cost of transmitting the respective additional data elements.

### 4.1 Parallel Traversal of the Reduction Tree

We implement the post-order traversal of the reduction tree via a stack *S* (Figure 3.a). Again, let *E* = (*e*_0_, *e*_1_, …, *e*_*n−*1_) be the *ordered* sequence of elements to be reduced. In the reduction tree, each *leaf* (in-degree 0) corresponds to an element *e*_*i*_ *∈ E*. When we encounter a leaf during the post-order traversal, we push the corresponding element onto *S*. We denote this as *addValue*. Conversely, each *non-leaf* node of the reduction tree represents an intermediate result *r*_*j*_, which we index in post-order. When we reach a non-leaf node, we *pop* the two topmost elements from *S*, apply the reduction operator ⊕, and *push* the result back onto *S*. We denote this as *reduceTwoValues*. Ultimately, the root of the reduction tree (out-degree 0) contains the final result of the reduction. This will be the sole remaining element on the stack. Therefore, we can implicitly represent the reduction via a sequence of *addValue* and *reduceTwoValues* operations.

**Fig. 1.**
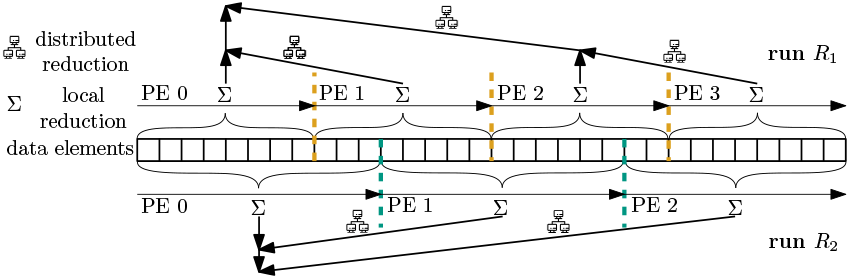
Common distributed-memory parallel reduction algorithms are non-reproducible if the PE-count differs between runs. First, each PE reduces its local elements to a single, intermediate, per-PE result. Next, these intermediate results are exchanged via messages over the network and further reduced into a single element. Here, the PE-count influences the reduction operation order and thus how results are rounded.

**Fig. 2.**
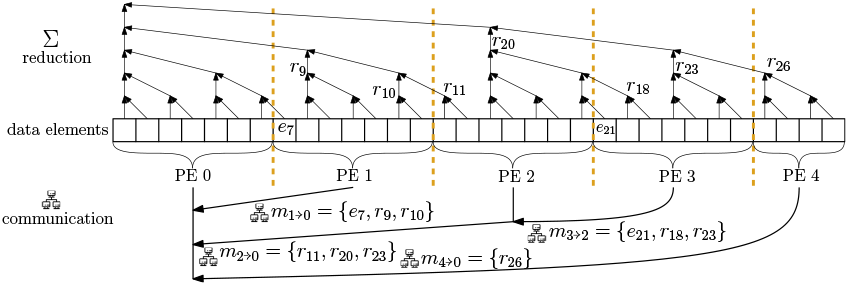
Decoupling the reduction operation order from the communication pattern. ReproRed reduces the data elements along a fixed tree, regardless of the number and topology of the PEs. The *p* PEs communicate via a binomial tree, which requires exchanging *p −* 1 messages (the theoretical minimum).

**Fig. 3.**
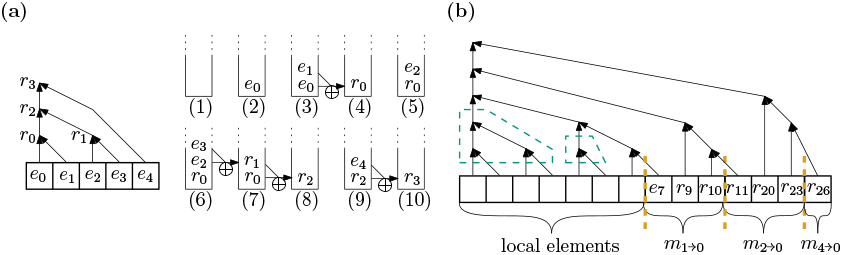
Implementation of ReproRed. **(a)** A post-order traversal of a binary tree (left) can be implemented via a stack (right). When we encounter a leaf node during the traversal, we push the respective data element onto the stack. For a non-leaf node, we pop the two topmost elements from the stack, reduce them, and push the result back onto the stack. At the root of the tree, the remaining stack element contains the final result of the post-order traversal. **(b)** Reduction operations performed on PE 1 in Figure 2. The received messages *m*_*i→j*_ are stored consecutively after the local elements. Note that some received elements already constitute intermediate results of the overall reduction. Further, the PE can reduce most of its local elements (green, dashed boxes) and thereby overlap computation with communication.

To parallelize this traversal – and thus the reduction – we distribute this sequence of *addValue* and *reduceTwoValues* operations across all PEs. For this, we assign each *addValue* operation to the PE that holds the respective data element. Each *reduceTwoValues* operation corresponds to a specific non-leaf node in the reduction tree. This node, including its descendants, induces a subtree in the reduction tree, with a set of (consecutive) input data elements as leaf nodes. Each of these elements resides on a specific PE. The PE set on which the elements of a reduction subtree reside have a joint Lowest Common Ancestor (LCA) in the communication tree. We assign the *reduceTwoValues* operation to this LCA-PE.

We determine this assignment of operations to PEs via a pre-processing step, which we execute once for each data distribution. Thus, when one invokes the distributed reduce operation with concrete values – possibly thousands of times per second (Section 6.5) – each PE already knows all operations it needs to execute *a priori*. (Figure 3.b).

During a reduction, each PE initially triggers the asynchronous message-receive to overlap computation with communication. As each PE knows the number of messages and data elements it will receive, it can arrange local and received data at consecutive memory locations. The operations of a PE correspond to pair-wise reductions in a subtree of the overall reduction tree. This subtree has input data elements and intermediate results computed on other PEs as leaves. When reducing this local subtree, we apply a fast base-case for fully local data; for example, we parallelize summation via manually-optimized AVX2 code (see Section 4.2). If a PE needs to perform a *reduceTwoValues* operation for which one operand has not been received yet, the PE blocks until the respective message exchange has been completed. Upon completing its reductions, the PE sends the computed intermediate results to its communication tree parent. Note that the sending PE’s stack will already contain the elements in the order that is expected by the receiving PE.

### 4.2 Applied Optimizations

ReproRed overlaps computation with communication (Figure 3) by utilizing non-blocking MPI messages, and thereby accelerates its runtime by 4.4% (median; data not shown). Further, ReproRed can reduce PE-local elements via a fast base-case, as long as the operation order remains unaltered. We provide respective vectorized kernels for summation, which ReproRed recursively applies to reduce four (SSE3) or eight (AVX2) elements or intermediate results (supplement). Our manually-optimized AVX2 kernel exhibits a speedup of 2.9% (median) over the compiler-vectorized reference on the datasets in Section 6.4. For datasets with tens of millions of elements across all PEs, we attain speedups of up to 200%. Moreover, we communicate via a *B*^4^, instead of a *B*^2^ tree (Section 2.1; not shown in all figures). We experimentally determined this degree. It attains a speedup of 5.6% (median; data not shown) over a *B*^2^ tree. This is due to a larger fraction of values being reduced at lower levels of the communication tree, which reduces message sizes by decreasing the number of times a single element is forwarded.

## 5. Reproducible Phylogenetic Inference

We present Repro-RAxML-NG, a proof-of-concept bit-reproducible version of the widely used phylogenetic tree inference tool RAxML-NG (developed by our lab). Repro-RAxML-NG currently supports bit-reproducible tree evaluation and tree searches. At present, the pattern compression, tip inner, and site repeats optimizations as well as the vectorization of phylogenetic likelihood derivatives required for branch length optimizations, are disabled to keep code changes to a necessary minimum for our experiments. Disabling these optimizations caused a slowdown of 88 *±* 38% (gemetric mean *±* standard deviation; see supplement). We experimentally verify that Repro-RAxML-NG yields bit-identical results under varying PE-counts on the empirical datasets described in Section 6 – some of which are known to cause diverging tree searches.

## 6. Evaluation

We conduct our experiments on the SuperMUC-NG HPC system.^1^ Each compute node comprises 96 GiB of memory and two Intel Skylake Xeon Platinum 8174 processors with 24 cores, each running at a fixed 2.3 GHz. The nodes communicate via an OmniPath network with a fat tree topology and a bandwidth of 100 Gbit s^*−*1^. The operating system is SUSE Linux Enterprise Server 15 SP3 running Linux Kernel version 5.3.18-150300.59.63. We compile our benchmark applications using GCC version 12.2.0 with full optimizations enabled (−O3) and all assertions in RAxML-NG disabled; we employ IntelMPI 2021.9.0. Repro-RAxML-NG^2^ is based on RAxML-NG commit 816f9d13^3^, which we use as a reference.

We perform reproducibility experiments on 10 130 empirical phylogenetic datasets from TreeBASE [26], comprising more than 100 000 distinct taxa collected from over 4000 publications. These TreeBASE datasets contain 4 to 4019 taxa, 1 to 179 partitions, and 21 to 3 848 295 sites – with up to 214 492 unique columns in the respective MSAs. For our runtime performance benchmarks, we utilize the same datasets previously used to evaluate the performance of RAxML-NG upon its release [21]. These datasets comprise between 1371 and 21 410 970 sites, and some are thus suitable for parallel inference on dozens or hundreds of PEs.

### 6.1 Reproducibility of Phylogenetic Inference

For each TreeBASE [26] dataset, we execute the same RAxML-NG [21] application binary eight times, using identical hardware, search settings, and random seeds. For each of the eight runs we exclusively vary the degree of parallelization. Specifically, we conduct inferences using one to five PEs with AVX2 vectorization, as well as sequential (single core) inferences with SSE3, AVX, AVX2, and auto-vectorization.

We quantify the divergence between the resulting trees using the relative Robinson-Foulds distance [27], which corresponds to the proportion of non-trivial bipartitions (i.e., bipartition induced by cutting *inner* branches) that differ between the two phylogenetic trees. The set of non-trivial bipartitions induced by all inner edges fully describe a tree.

When multiple trees exhibit sufficiently similar likelihood scores, there is no compelling reason to favor one over the others. Thus, following Shen *et al*. [30], we apply the Approximately Unbiased (AU) test [31] to determine if two topological distinct trees – resulting from inferences under different levels of parallelism – exhibit significantly different likelihood scores (*p <* 0.05). To prevent erroneous rejection of the correct tree in sets with similar likelihoods [31], we consider trees with a log-likelihood difference of less than 10^*−*3^ to not be significantly different.

### 6.2 Results

We consider a dataset as *diverging* if the eight phylogenetic inferences conducted on it yield at least two topologically distinct trees. In our experiments, 31% (3111) datasets are diverging (Figure 4). Further, 46% (1418) of these diverging datasets exhibit a log-likelihood difference exceeding 10^*−*3^ log-likelihood units, and 8% (245) contain significantly different trees (AU-test, *p <* 0.05) in the resulting tree set that comprises a total of eight trees (Figure 5).

**Fig. 4.**
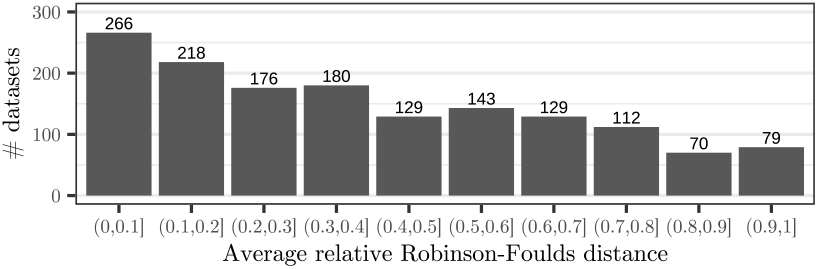
Average relative Robinson-Foulds distance between all distinct tree topologies resulting from tree searches that diverged due to different parallelization degrees. We only show datasets that yielded at least two (out of eight) distinct tree topologies (31% of all datasets).

**Fig. 5.**
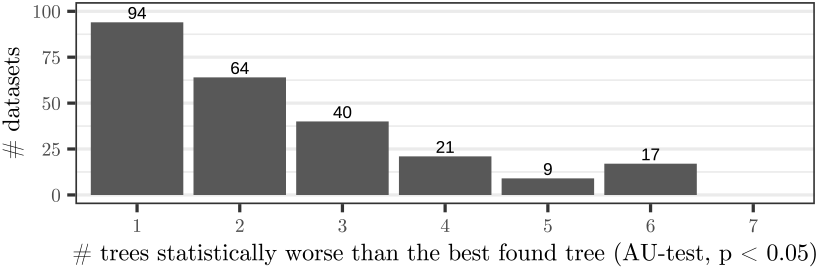
Number of trees resulting from a diverging tree search that are significantly worse than the best-known tree. We only show the 8% of diverging datasets where the tree searches yielded at least one tree that is significantly worse than the best-known tree (AU-test, *p <* 0.05).

Compared to Shen *et al*. [29], who analyzed 3515 single-gene datasets, we observe a higher proportion of diverging datasets (31% vs. 9.3% with RAxML-NG), yet fewer significant differences (8% vs. 25% of diverged datasets). This is likely due to the fact that we assess eight distinct parallelization degrees, instead of just two, as in Shen *et al*. [29]. Further, we classify trees with a LLH-difference *<* 10^*−*3^ as *not being statistically different*, while Shen *et al*. [29] do not apply this (arbitrary) threshold. Without this filtering step, 38% (1186) of our diverging datasets yield significantly different trees (AU-test, *p <* 0.05).

### 6.3 Conclusion

Supporting Shen *et al*. [30], we demonstrate that phylogenetic tree searches are sensitive to different parallelization degrees as these cause tree searches to diverge because of distinct rounding errors. This divergence introduces an unknown bias in the distribution of the resulting trees. This is orthogonal to the tree differences caused by variations in random seeds or bootstrapping [12], which are well studied [14]. Whether differences between trees that are considered as being not significantly different from each other also have implications on their biological interpretation is non-trivial to answer and remains an open question. Therefore, we advocate for using bit-reproducible implementations of phylogenetic tree searches.

### 6.4 Runtime of Isolated Bit-Reproducible Reduction

We compare the runtime of the bit-reproducible *all-reduce* algorithms ReproBLAS (state-of-the-art), ReproRed (ours), and Gather-Bcast (recommended by the MPI-standard), as well as the non-reproducible `MPI Allreduce` and Reduce-Bcast (Section 3). As only ReproRed supports reduction operators beyond summation, we compare summation performance.

As the exact data distribution impacts ReproRed’s runtime, we extract realistic data distributions from our phylogenetic inferences (Section 5). Concretely, we distribute 1000 to 4 110 000 double precision (64 bit) IEEE 754 floating-point elements across 48 to 768 PEs, maintaining an approximately equal numbers of elements per PE. We perform 5000 iterations for each data distribution, discarding the first 1000 runs for each distribution to warm up the network caches. The absolute runtimes for `MPI Allreduce` range from 1.6 to 600 µs (median 10.8 µs). We verify that the reductions performed with ReproBLAS and ReproRed yield bit-identical results across different data distributions and PE-counts. Additionally, we verify that the results of all algorithms differ by less than 10^*−*6^. Gather-Bcast (Section 3) is 9.8 to 125 (median 31) times slower than `MPI Allreduce`. In addition, it requires at least one of the PEs to have sufficient memory to store *all* elements. We thus consider Gather-Bcast impractical and omit it in Figure 6. The ReproRed-TwoPhase *all-reduction* consists of a ReproRed reduction and a subsequent `MPI Bcast`. To evaluate the overhead induced by this two-phase strategy, we measure the runtime of a non-reproducible local reduction followed by an `MPI Reduce` and an `MPI Bcast` (*Reduce-Bcast*; Figure 6). The median number of elements per PE in our phylogenetic experiments is 1935, yielding Reduce-Bcast 61% slower (median) than all-reduce. However, for more than 45 000 elements per PE, Reduce-Bcast is merely 5% slower (median). We find that ReproRed outperforms ReproBLAS-TwoPhase under most data distributions exceeding 3000 elements per PE, and outperforms ReproBLAS-Allreduce exceeding 10 000 elements per PE (Figure 6). Both, ReproRed and ReproBLAS exchange only the theoretical minimum number of messages for a reduction. While ReproRed performs less local work than ReproBLAS, it also increases the number of computations along the critical path (supplement). In accordance to this, we observe that ReproRed is slower than ReproBLAS for less than a few thousand elements per PE. However, as the number of elements per PE increases, local computation starts compensating these effects. This explains ReproRed’s notably faster performance on data distributions with tens of thousands of elements per PE (Figure 6). For instance, ReproRed is 37% (median) times faster than ReproBLAS on 8000 to 12 000 elements per PE and 92% (median) times faster on more than 30 000 elements. Note, however, that ReproBLAS utilizes additional local work to guarantee the final result accuracy. In contrast, ReproRed’s pair-wise summation merely increases the accuracy [15]. Further, ReproBLAS is restricted to summation as reduction operator, while ReproRed supports arbitrary associative reduction operators.

**Fig. 6.**
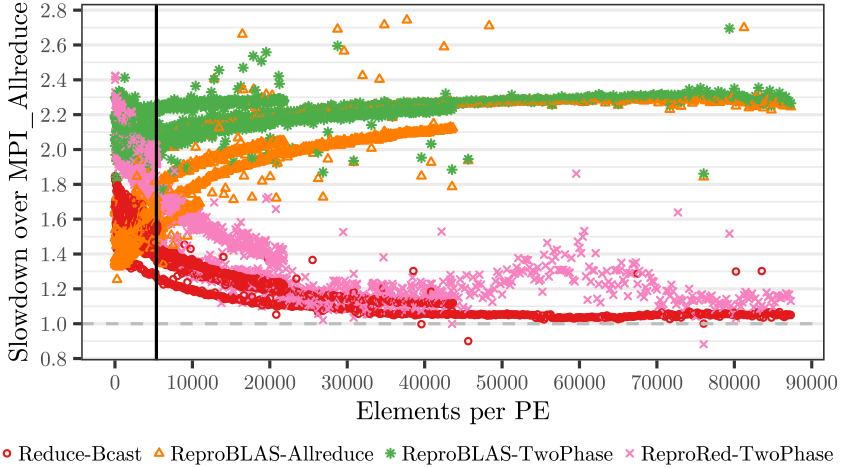
Slowdown of bit-reproducible all-reduce algorithms over a non-reproducible all-reduction. Each point represents the median runtime of 4000 iterations for the specific number of elements and number of PEs combination divided by the respective median runtime of the non-reproducible reference (`MPI Allreduce`). The vertical line represents the median number of per-PE log likelihood values in our empirical data analyses (Section 6.5).

### 6.5 Bit-Reproducible Phylogenetic Tree Search

We assess the slowdown of our bit-reproducible Repro-RAxML-NG (Section 5) in comparison to the unmodified reference RAxML-NG (Figure 7). Specifically, we evaluate Repro-RAxML-NG with the reproducible all-reduce algorithms ReproBLAS (state-of-the-art), Gather-Bcast (recommended by the MPI-standard), and ReproRed (ours). Further, we present results for the non-reproducible `MPI_Allreduce` and TwoPhase approaches. We use the PE counts recommended by RAxML-NG’s `--parse` for the respective datasets, resulting in 253 to 11 903 (median 5569) unique MSA columns per PE and 1935 (median) all-reduce operations per second (for the RAxML-NG reference version). For consistency, we disable optimization features not supporting bit-reproducibility in all variants (Section 5). We exclude datasets where the tree searches diverged, as this yields the runtimes not comparable and verify that the resulting log likelihood scores of trees differ by less than 10^*−*3^ log likelihood units among all variants.

**Fig. 7.**
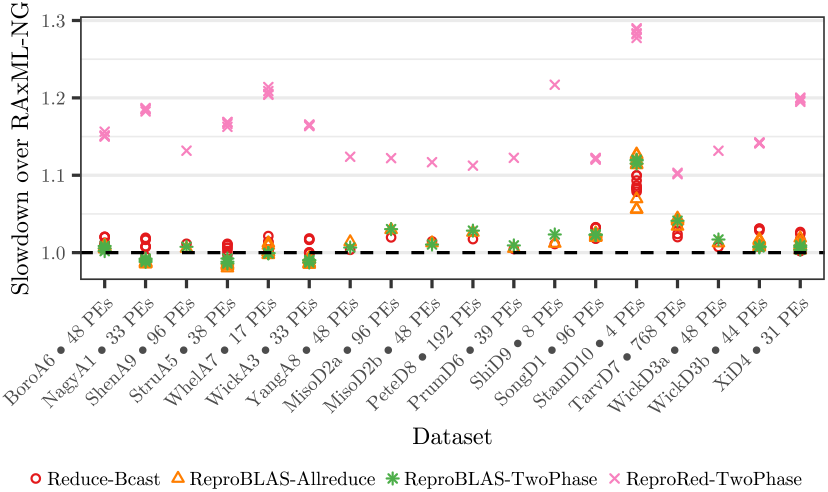
Slowdown of bit-reproducible Repro-RAxML-NG over reference RAxML-NG. Reduce-Bcast uses a non-reproducible reduction but includes the remaining modifications required for bit-reproducibility. The variants ReproBLAS and ReproRed yield bit-reproducible results.

Repro-RAxML-NG using the Gather-Bcast all-reduce algorithm is 21 to 386% (median 26%) slower than the reference. It is consistently the slowest variant across all configurations; we hence omit it from further discussion.

Repro-RAxML-NG ReproRed-TwoPhase is 10 to 29% (median 17%) slower than the reference. However, when employing the bit-reproducible ReproBLAS all-reduction, reproducibility induces only 0 to 12.7% (median 0.8%) slowdown. This is consistent with the isolated measurements (Section 6.4), where ReproBLAS outperforms ReproRed-TwoPhase on less than 10 000 elements per PE. Even though some of RAxML-NG’s optimizations are disabled, due to lack of bit-reproducible implementations (Section 5), we are cautiously optimistic that implementing bit-reproducible phylogenetic tree inference tools is feasible.

## 7. Conclusion and Future Work

We highlight the issue of non-reproducibility in phylogenetic tree inference and present the first (proof of concept) bit-reproducible phylogenetic tree inference tool. We consider this an important step towards bit-reproducible – and thus more reliable – phylogenetic tree inferences. We also develop the first open-source bit-reproducible reduction algorithm supporting associative reduction operators, which we already integrated into the open-source MPI-wrapper library KaMPIng [34] to facilitate the transition to bit-reproducible code.

In future work, we aim to develop bit-reproducible kernels for calculating the derivatives of the phylogenetic likelihood and to port optimizations like pattern compression [33] to Repro-RAxML-NG. We also intend to implement a recursive-doubling all-reduce variant of ReproRed (see supplement), to further accelerate all-reduction. Additionally, we plan to extend ReproRed to support both, inclusive, and exclusive prefix sums.

## Supporting information

On-Line Supplement

## Footnotes

1 https://doku.lrz.de/display/PUBLIC/SuperMUC-NG

2 https://doi.org/10.5281/zenodo.15017407

3 https://github.com/amkozlov/raxml-ng/commit/816f9d13

