## Supplementary material for "Bit-Reproducible Phylogenetic Tree Inference under Varying Core-Counts via Reproducible Parallel Reduction Operators": On-Line Supplement

#### 1 Binomial Trees

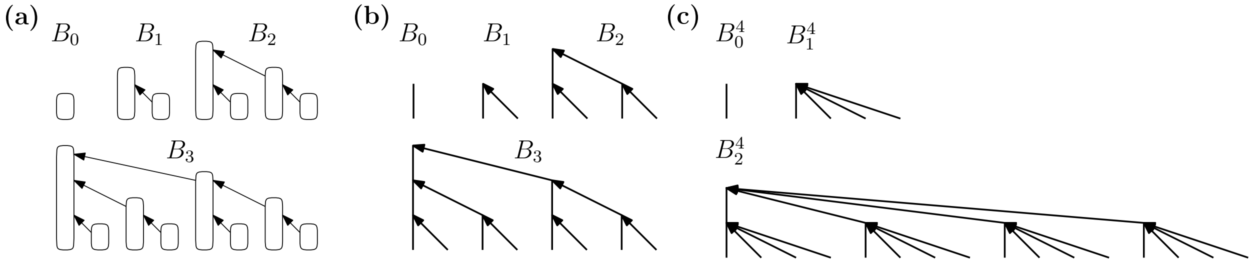


**Figure 1: (a)** (Canonical) binomial trees $B_0$ to $B_3$. **(b)** Alternative representation of $B_0$ to $B_3$, indicating the messages sent in a binomial tree reduction. **(c)** $B^4$ binomial trees.

Following Cormen *et al.* [2, ch. 19.1.1], we define a binomial tree as follows: The binomial tree $B_0$ of order 0 consists of a single node. A binomial tree $B_k$ of order $k$ comprises a root node whose children are the roots of the binomial trees of order $k-1, k-2, \dots, 2, 1, 0$ (Figure 1.a). Further, we define a generalized binomial tree $B^m$, where the root of a binomial tree $B^m_k$ has $m-1$ copies of each $B^m_i$ for $i = k-1, k-2, \dots, 2, 1, 0$ as children (Figure 1.b). Thus, in this notation, a canonical/ binomial tree is a $B^2$.

In a binomial tree reduction, PEs send intermediate results along the edges of a binomial tree towards the root and also reduce the values in each step. For a small number of elements per PE, a binomial tree reduction is optimal in terms of message-count and size, as well as the number of operations performed, and thus, also running time [8, ch. 13.1.1].

#### 2 Optimizations applied to ReproRed

The three key optimizations we implement in ReproRed are to overlap computation with communication, employing a fast base-case for reducing PE-local data elements, and to communicate via a $B^4$ instead of a $B^2$ binomial reduction tree (Figure 1).

**Vectorized Reduction of PE-local Subtrees** We implement AVX2 (Figure 2) and SSE3 (not shown) kernels for vectorizing the base case of summation as the reduction functions. Importantly, our binary reduction tree allows for parallelizing the reduction without altering the order of operations. In contrast, a left-to-right reduction is inherently sequential and requires vectorized algorithms to change the operation order, thereby compromising bit-reproducibility. While we implement an AVX2-vectorized base-case for *summing* over PE-local subtrees in the reduction tree, we expect analogous fast base-case implementations to be feasible for numerous reduction operations.

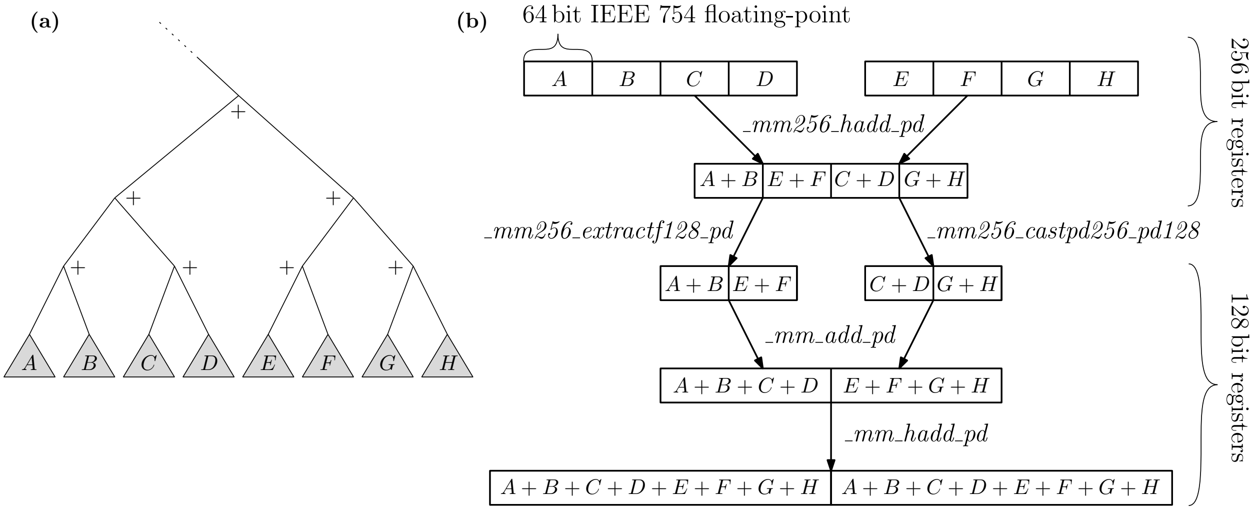


**Figure 2:** Pair-wise addition of eight 64 bit IEEE 754 floating-point values using AVX2 vectorization. **(a)** The subtree of the reduction tree to be reduced. $A, \ldots, H$ can be either input data elements or intermediate results from lower subtrees. **(b)** Implementation of the AVX2-vectorized reduction of the subtree depicted in (a). Note that the operation order is consistent with a non-vectorized pair-wise addition implemented as a post-order traversal as described in the main text.

Our AVX2 implementation reduces a subtree comprising eight elements or intermediate results using two 256 bit registers, two 128 bit registers, two load operations, and five additional SIMD instructions (Figure 2.b). We apply the reduction recursively, that is, the leaves of this eight-element tree may be intermediate results of preceding vectorized reductions. For example, to reduce 512 input elements, we execute three levels of AVX2-optimized reductions, resulting in 73 invocations of the 8-value AVX2 kernel.

First, we horizontally sum pairs of elements in two 256 bit registers and combine the results into a single 256 bit register (Figure 2.b). Since AVX2 instructions operate within 128 bit lanes, we must separately extract the upper and lower 128 bit segments after the initial horizontal addition in order to maintain the correct summation order. Subsequently, we conduct an element-wise addition on the two resulting 128 bit registers, followed by horizontal addition on the resulting 128 bit register. Again, as the operation order and thus the round-off errors remain unaltered, a vectorization with different SIMD register widths will produce identical results.[1]

Compilers, such as GCC, attempt to automatically vectorize code.[2] However, a reduction with manually optimized AVX2 vectorization achieved a speedup of 2.9 % (median) compared to the compiler-vectorized reference on the datasets described in the main text. For datasets with tens of millions of elements across all PEs, we attain speedups of up to 200 %.

**Using a $B^4$ Reduction Tree** We implement the communication tree as a $B^4$, instead of a $B^2$ tree (Figure 1, not shown in all figures). We empirically determined this appropriate degree of $m = 4$ and gain a speedup of 5.6 % (median; data not shown) over using a $B^2$ tree. This acceleration is attained because ReproRed is thus able to reduce data elements at lower levels of the communication tree, thereby reducing message sizes and decreasing the number of times an element will be forwarded (Section 3).

**Overlapping Communication with Computation** We implement ReproRed such as to overlap computation with communication by utilizing non-blocking MPI messages. Specifically, each PE first posts the

[1]We experimentally verified that a SSE3 implementations yields bit-identical results with our AVX2 implementation.

[2]We manually confirmed that the relevant code sections compile to AVX machine instructions when providing the `-mavx` flag.

MPI receive operations before reducing its local elements. A PE blocks and waits for a message only when it requires an element that has not yet been received. By overlapping computation with communication, we accelerate the reduction by 4.4 % (median; data not shown).

#### 2.1 Bit-Reproducible All-Reduction using Recursive Doubling

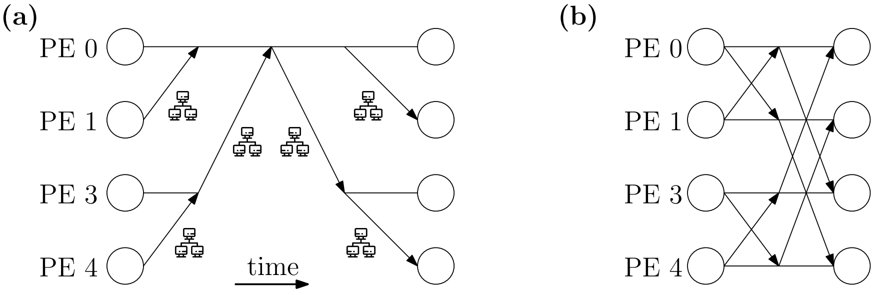


**Figure 3:** **(a)** The ReproRed-TwoPhase all-reduction algorithm consists of a binomial tree ReproRed reduction, followed by a binomial tree broadcast. **(b)** The ReproRed-RecursiveDoubling all-reduction algorithm is based on the recursive doubling all-reduction [10]. It reduces the latency from $2\log_2(p)$ to $\log_2(p)$ messages for $p$ PEs compared to ReproRed-TwoPhase.

In ReproRed, all-reduction is currently implemented as a binomial tree reduction followed by a (binomial tree) broadcast (Figure 3.a). This induces a latency of $2\log_2(p)$ messages for $p$ PEs. Here, we propose a variant with a latency of $\log_2(p)$ messages using the recursive doubling algorithm [10]. In the first step of recursive doubling, PEs with a distance of one exchange data. In a second step, PEs with a distance of two exchange data – including the data they received from the other PE in the first step. In subsequent steps, the distance increases exponentially, until, after $\log_2(p)$ steps, all PEs know each other PEs' data – or a reduced form thereof. Note that in recursive doubling, a PE always has knowledge of elements that reside on PEs that are *consecutive* in PE-id (*rank*)-space, thus allowing for the reduction of intermediate results (Figure 3.b). Future work will include implementing this ReproRed-RecursiveDoubling algorithm.

### 3 Factors Impacting ReproRed's Runtime

In the following, we discuss the factors impacting the runtime of ReproRed compared to ReproBLAS: The number and size of messages exchanged, the local work, as well as the length of the critical path.

#### 3.1 Message Size And Count

For a *reduction*, each of the $p$ PEs, except the root, has to send at least one message in order for its data elements to be included in the end result. This yields the lower bound of $p-1$ on the *message count* [8, ch. 13.9]. ReproRed sends exactly $p-1$ messages along the edges of a binomial tree. This also holds for the $B^4$ tree, as again, each PE, except the root, sends exactly one message. ReproBLAS can utilize an arbitrary communication pattern, for instance, a binomial tree, which exchanges $p-1$ messages. Further, for an *all-reduction*, the two-phase approaches ReproRed-TwoPhase and ReproBLAS-TwoPhase (as well as Reduce-Bcast) exchange $2(p-1)$ messages. We experimentally verify in our benchmarks that the implementations of these algorithms exchange exactly the described number of messages.

The *message sizes* exchanged also do not differ substantially between ReproBLAS and ReproRed. ReproBLAS exchanges six double precision floating-point values, that is, 384 bit, per message [1] – which we experimentally verify for our benchmarks using the Score-P profiler [4]. Conversely, the message size in ReproRed varies depending on the number of elements, the number of PEs, and the distribution of the elements among the PEs.

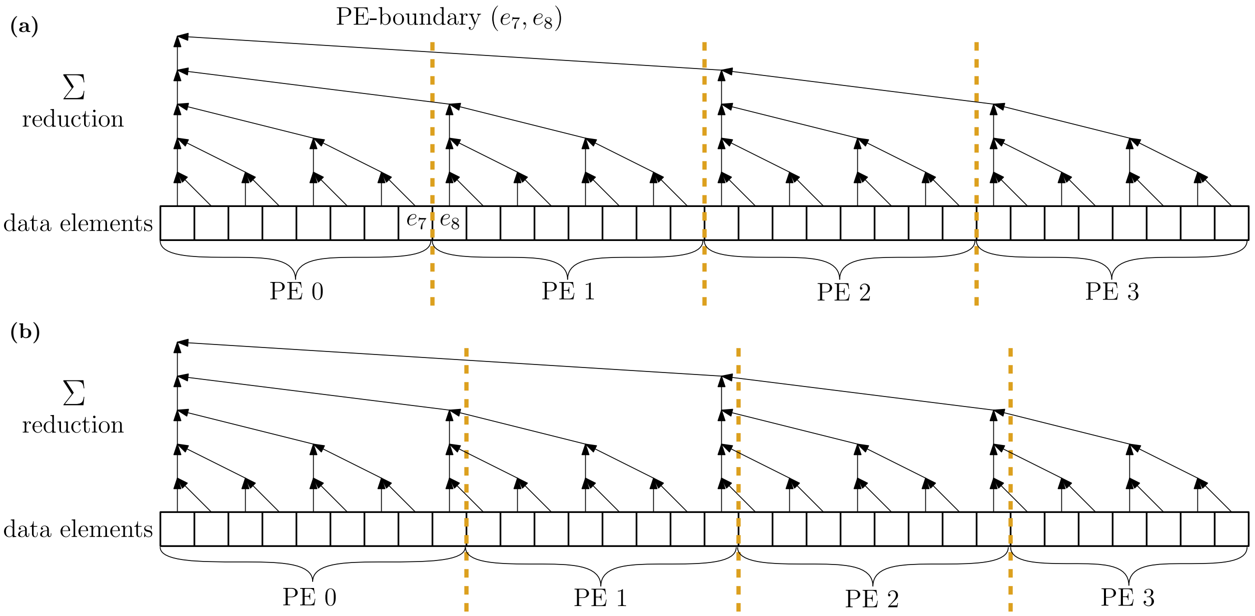


**Figure 4:** Examples of the best (a) and worst (b) case data distributions with regard to the message size (and critical path length) in ReproRed. Dashed lines represent PE-boundaries. **(a)** In the best case, each PE holds a power-of-two number of elements and is thus able to reduce its elements into a single intermediate result. Here, each PE-boundary cuts exactly one edge of the reduction tree. **(b)** In the worst case, the PE-boundaries cut $\mathcal{O}(\log_2(n))$ edges of the reduction tree. Each cut edge corresponds to an intermediate results that has to be exchanged over the network.

Consider, for instance, a distribution of $n$ elements $e_i$ ($i \in \mathbb{N}_0$) among $p$ PEs, such that each PE holds the same power-of-two number of elements (Figure 4.a). Specifically, PE $j$ holds elements $e_{j \cdot 2^k}, \ldots, e_{(j+1) \cdot 2^k - 1}$, where $k = log_2(n/p)$. We denote a pair $(e_i, e_{i+1})$ as a *PE-boundary* if $e_i$ and $e_{i+1}$ are held by different PEs. Under this data distribution, PE-boundaries are located in-between full subtrees of the binary reduction tree. Thus, each PE is able to reduce all elements it holds into a single intermediate result. Further, during communication, the PE can immediately reduce the incoming values with its local intermediate results. Consequently, each message has a length of exactly one intermediate result (64 bit).

In contrast, consider the distribution in Figure 4.b: The first PE holds one element more than a power-of-two, while all other PE still hold a power-of-two elements.[3] Specifically, PE 0 holds $e_0, e_{2^k+1}$ and a PE $j > 1$ holds the elements $e_{j \cdot 2^k + 1}, \ldots, e_{(j+1) \cdot 2^k}$. Under this distribution, the PE-boundaries cut $\log_2(n)$ edges of the binary reduction tree. Each cut edge represents an element or intermediate result that must be exchanged across the network. Moreover, the two PEs of a boundary might not share an edge in the communication tree. In such cases, the corresponding data must be sent up the communication tree until they reach the lowest common ancestor of the two PEs at the boundary.

In summary, each PE-boundary cuts up to $\log_2(n)$ edges of the reduction tree, resulting in the respective PE to send this number of elements to its parent-PE in the communication tree. Further, each of these elements is relayed up to $\log_2(p)$ times. We observe that the messages exchanged in our isolated benchmarks contain 1 up to 24 (median and mean 11) elements, each comprising 64 bit. Thus, we expect the startup overhead for each message to substantially exceed the cost of transferring these additional bytes [8, ch. 13.6.2].

[3] The last PE has one element less, which we omit in the following discussion for simplification.

#### 3.2 Local Work

ReproBLAS executes 9 floating-point and 3 bit-wise operations per floating-point element in the input, regardless of the data distribution [1]. In contrast, ReproRed requires five floating-point SIMD operations to sum eight fully-local floating-point values. Further, for each pair-wise reduction involving received data, ReproRed must increment an iterator, extract a bit from a bit-vector, and perform a conditional jump in order to decode the operation. Additionally, for each non-local reduction (`reduceTwoValues`) operation, ReproRed performs two `pop` and one `push` operation on a stack.

Using the Score-P profiler [4], we experimentally obtain that ReproBLAS requires more runtime for its local work compared to ReproRed in our isolated benchmarks. The precise overhead for local work of ReproBLAS over ReproRed depends on the number of elements per PE and varies by up to a factor of two (data not shown).

#### 3.3 Length of Critical Path During Message Exchange

We define an algorithm's *critical path* as the longest task sequence where each task depends on the preceding one. These tasks include message exchanges and local operations, such as applying a reduction operation to two elements. For both, ReproRed and ReproBLAS, the number of messages on the critical path in a *reduction* corresponds to the longest path in a binomial tree, that is $\log_2(p)$ [10], where $p$ is the number of PEs. ReproBLAS allows the PEs to reduce *all* their local elements into a single intermediate result in parallel. These elements are subsequently reduced via at least $\log_2(p)$ steps, using, for example, a binomial tree reduction. In each step, ReproBLAS executes 9 floating-point operations and 3 bit-operations [1]. Thus, ReproBLAS executes $\mathcal{O}(n/p + \log_2 p)$ operations on its critical path. In ReproRed, depending on the data distribution, PEs cannot reduce their local elements into a single intermediate results, but rather a (small) set of intermediate results. As each PE-boundary cuts up to $\log_2(n)$ edges of the reduction tree, the respective number of operations is thus located on the critical path. Overall, assuming $n > p$, there are $\mathcal{O}(n/p + \log_2(n))$ pair-wise reductions – and thus arithmetic operations – on ReproRed's critical path. Note that the critical paths are of the same length asymptotically, but differ in a constant factor. We implement ReproRed to overlap communication with computation, thereby mitigating the impact of this extended critical path (Section 2).

During an *all-reduction*, the two-phase approaches, ReproRed-TwoPhase, ReproBLAS-TwoPhase, and Reduce-Bcast, perform a binomial tree reduction followed by a binomial tree broadcast. Thus, these methods exchange $2\log_2(p)$ messages on the critical path. However, all-reduce algorithms with only $\log_2(p)$ messages on their critical path exists, such as recursive doubling [10]. We outline a recursive doubling based implementation of ReproRed in Section 2.1. Thus, when assessing ReproRed's *reduction* performance, one can argue for using Reduce-Bcast as the reference benchmark.

### 4 Datasets for Repro-RAxML-NG Benchmarks

For our end-to-end Repro-RAxML-NG experiments, we employ the same datasets used to benchmark RAxML-NG's performance upon its release [6]. These datasets are described in the original paper's online supplement[4] (Table 3) and available on Figshare[5].

[4]`https://academic.oup.com/bioinformatics/article/35/21/4453/5487384#supplementary-data`
[5]`https://figshare.com/s/6123932e0a43280095ef?file=13639241`

### 5 Reproducing our Runtime Benchmark Results

We provide an automated setup to reproduce our isolated all-reduce and Repro-RAxML-NG[6]. To ensure that all executions of the benchmark utilize an identical binary and shared libraries[7], the script downloads a container image and runs it via Charliecloud [7] – container solution that does not require special privileges and is therefore suited for high-performance computing contexts. Please refer to the `README` of the linked benchmark repository for further information on how to reproduce our results.

### 6 Verification of Bit-Reproducibility

To verify the reproducibility of ReproRed, we ran all three summation algorithms with different core-counts in steps of 48 ($p \in \{48, \ldots, 768\}$) on randomly generated data with the same length as our benchmark datasets. With ReproBLAS and ReproRed, we detected no deviation between results for all test datasets, while values produced by all-reduce were already irreproducible between runs with $p = 48$ and $p = 96$ PEs. Table 1 shows the difference between the largest and smallest result collected over all values of $p$. For ReproBLAS and ReproRed, this value is zero,[8] indicating that the result is independent of the number of PEs. For all-reduce, the largest observed relative error was $7.45 * 10^{-9}$ (dataset TarvD7), which could potentially cause tools like RAxML-NG to dismiss certain trees during likelihood maximization leading to diverging tree searches. Results from ReproBLAS and ReproRed were also reproducible across different machines and compiler versions.[9]

**Table 1:** Difference between smallest and largest obtained sum from runs with varying PE-count.

| **Dataset** | $\Delta$ `std::accumulate` + **AllReduce** | $\Delta$ **ReproBLAS** | $\Delta$ **ReproRed** |
|---|---|---|---|
| BoroA6 | $8.73 * 10^{-11}$ | 0 | 0 |
| ChenA4 | $4.65 * 10^{-10}$ | 0 | 0 |
| KatzA10 | $3.63 * 10^{-12}$ | 0 | 0 |
| MisoA2 | $8.73 * 10^{-11}$ | 0 | 0 |
| MisoD2a | $1.16 * 10^{-10}$ | 0 | 0 |
| MisoD2b | $2.91 * 10^{-11}$ | 0 | 0 |
| NagyA1 | $1.45 * 10^{-11}$ | 0 | 0 |
| PeteD8 | $1.39 * 10^{-9}$ | 0 | 0 |
| PrumD6 | $5.82 * 10^{-11}$ | 0 | 0 |
| ShenA9 | $5.82 * 10^{-11}$ | 0 | 0 |
| ShiD9 | $3.63 * 10^{-12}$ | 0 | 0 |
| SongD1 | $3.49 * 10^{-10}$ | 0 | 0 |
| StamD10 | $1.13 * 10^{-13}$ | 0 | 0 |
| StruA5 | $4.36 * 10^{-11}$ | 0 | 0 |
| TarvD7 | $7.45 * 10^{-9}$ | 0 | 0 |
| WhelA7 | $3.63 * 10^{-12}$ | 0 | 0 |
| WickA3 | $1.45 * 10^{-11}$ | 0 | 0 |
| WickD3a | $5.82 * 10^{-11}$ | 0 | 0 |
| WickD3b | $5.82 * 10^{-11}$ | 0 | 0 |
| XiD4 | $2.91 * 10^{-11}$ | 0 | 0 |
| YangA8 | $5.82 * 10^{-11}$ | 0 | 0 |

[6] `https://github.com/stelzch/reproducible-reduce-benchmark`

[7] Note that this script is newer than the benchmarks described in the main text, and we thus expect the LLH-values to differ slightly. We refrain from re-running our experiments in order to conserve computational end ecological resources.

[8] Or at least smaller than the machine epsilon for IEEE 754 double precision floating-point numbers.

[9] Linux 5.4.0-89-generic with GCC 9.4.0 on i10pc138, Linux 4.18.0-193.65.2.el8_2.x86_64 with GCC 11.2 on bwUniCluster 2.0

### 7 Slowdown Caused by Disabling RAxML-NG's optimizations

We present Repro-RAxML-NG, a proof-of-concept bit-reproducible version of the widely used phylogenetic tree inference tool RAxML-NG (developed by our lab). Repro-RAxML-NG currently supports bit-reproducible tree evaluation and tree searches. At present, the pattern compression [9], tip inner, and site repeats [5] optimizations as well as the vectorization of phylogenetic likelihood derivatives [3] required for branch length optimizations, are disabled to keep code changes to a necessary minimum for our experiments. Disabling these optimizations caused a slowdown of $88 \pm 38\,\%$ (geometric mean $\pm$ standard deviation; Table 2). These simplifications do not affect reproducibility for we plan to reintroduce them to Repro-RAxML-NG.

**Table 2:** Slowdown Caused by Disabling RAxML-NG's pattern compression, tip inner, and site repeats optimizations as well as the vectorization of phylogenetic likelihood derivatives required for branch length optimizations.

| **Dataset** | **Slowdown** |
|---|---:|
| BoroA6 | 1.55 |
| ChenA4 | 1.63 |
| KatzA10 | 1.94 |
| MisoA2 | 1.69 |
| NagyA1 | 1.44 |
| ShenA9 | 1.61 |
| StruA5 | 1.56 |
| WhelA7 | 1.50 |
| WickA3 | 1.53 |
| YangA8 | 1.72 |
| MisoD2a | 1.85 |
| MisoD2b | 2.23 |
| PeteD8 | 2.10 |
| PrumD6 | 2.31 |
| ShiD9 | 2.73 |
| SongD1 | 2.15 |
| StamD10 | 2.10 |
| TarvD7 | 1.98 |
| WickD3a | 2.10 |
| WickD3b | 2.26 |
| XiD4 | 2.01 |
